# Microstructured silk-fiber scaffolds with enhanced stretchability

**DOI:** 10.1101/2023.06.01.542724

**Authors:** Martina Viola, Gerardo Cedillo-Servin, Anne Metje van Genderen, Isabelle Imhof, Paula Vena, Marko Mihajlovic, Susanna Piluso, Jos Malda, Tina Vermonden, Miguel Castilho

## Abstract

Despite extensive research, current methods for creating three-dimensional (3D) silk fibroin (SF) scaffolds lack control over molecular rearrangement, particularly in the formation of β-sheet nanocrystals, as well as hierarchical fiber organization at both micro- and macroscale. In this study, we introduce a fabrication process based on electrowriting of aqueous SF-based solutions followed by post-processing using an aqueous solution of sodium dihydrogen phosphate (NaH_2_PO_4_). This approach enables hierarchical assembly of SF chains via β-sheet and α-helix formation. Moreover, this process allows for precise control over micro- and macro-architectures in microfiber scaffolds, enabling the creation of 3D flat and tubular macrogeometries with square-based and crosshatch microarchitectures, featuring inter-fiber distances of 400 µm and approximately 97% open porosity. Remarkably, the printed structures demonstrated restored β-sheet and α-helix structures, which imparted an elastic response of up to 20% deformation and the ability to support cyclic loading without plastic deformation. Furthermore, the printed constructs supported *in vitro* adherence and growth of human conditionally immortalized proximal tubular epithelial cells and glomerular endothelial cells, with cell viability above 95%. These cells formed uniform, aligned monolayers that deposited their own extracellular matrix. These findings represent a significant development in fabricating organized SF scaffolds with unique fiber structures, mechanical and biological properties, making them highly promising for regenerative medicine applications.

## 1. Introduction

Polymers, particularly those derived from natural sources, are becoming increasingly popular as bio-interactive materials for regenerative medicine (RM). Among these, silk fibroin (SF), a natural polymer obtained from silk, stands out as a highly attractive material due to its biocompatibility, biodegradability, and exceptional mechanical properties.^1, 2^ These properties stem from SF’s unique chemical composition and hierarchical organization at different length scales. At the nanoscale, SF is composed of β-sheet nanocrystals embedded in a softer semi-amorphous phase consisting of random coil chains. These nanostructures further assemble into fibrils at the microscale, which then bundle into fibers or threads at the macroscale (Figure 1.1).^3, 4^ Although the assembly process is not fully understood, it is thought to occur during silk fiber spinning by spiders or worms, through the formation of a micellar-like pre-assembled silk protein that solidifies upon ejection from the spinning duct.^5^ This process takes place under ambient conditions and is accompanied by acidification of the surrounding environment, water loss, gradients in salt ions (such as sodium and potassium) and exposure to shear and elongation forces,^6, 7^ ultimately resulting in the transition of silk micelles to water-insoluble silk fibers.^7^

**Figure 1.**
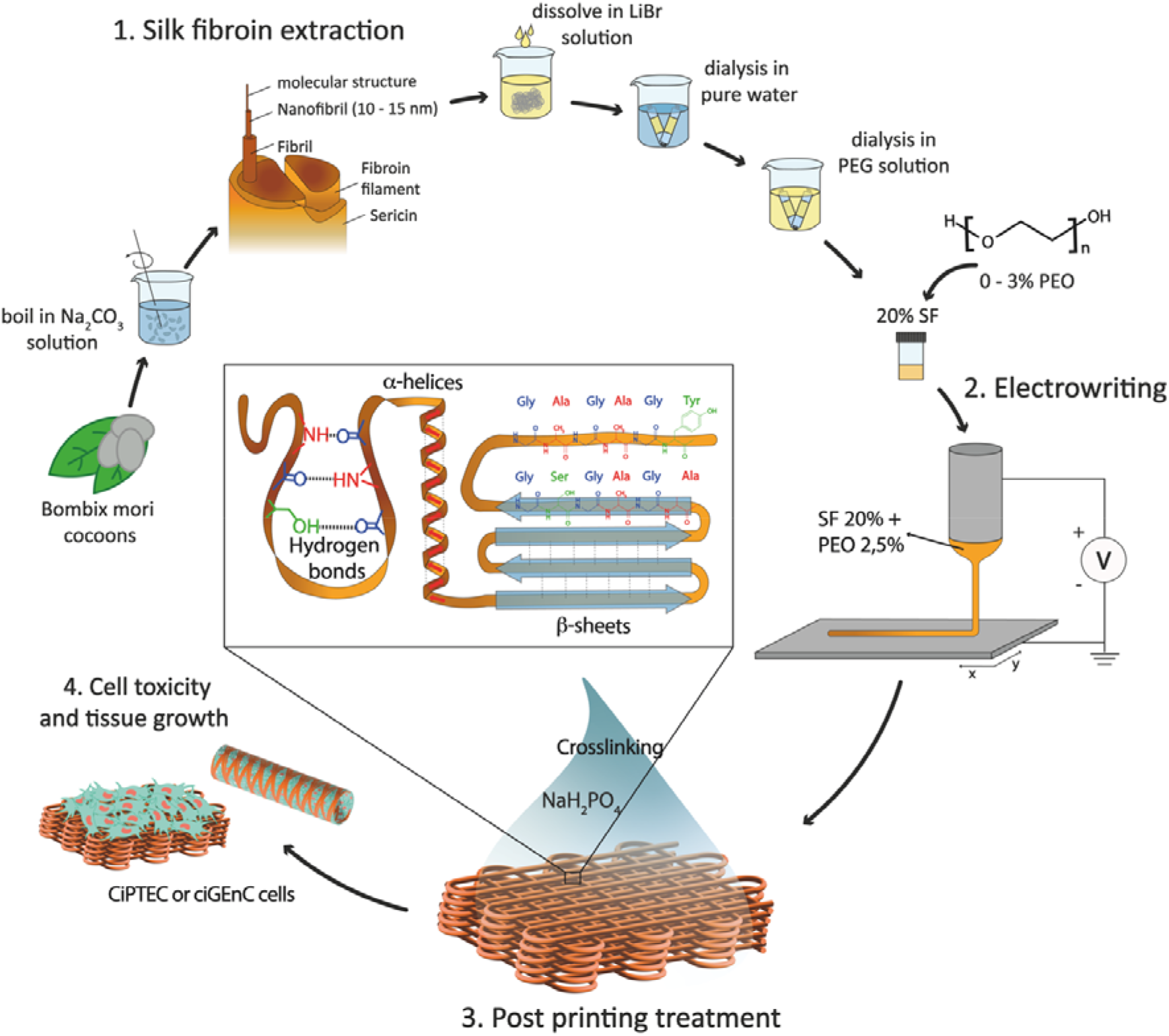
Schematic representation of aqueous electrowriting of organized SF fiber structures. 1. extraction of SF from *Bombix mori* cocoons and addition of PEO. 2. Electrowriting process of SF biomaterial inks, and 3. Post processing treatment with NaH_2_PO_4_, to induce secondary structure of the protein from random coil to β-sheets and α-helices. 4. Conditionally immortalized proximal tubule epithelial cells (ciPTEC) and conditionally immortalized glomerular endothelial cells (ciGEnC) toxicity and tissue growth on well-organized flat and tubular SF fiber scaffolds.

Current methods for creating biomimetic SF fibers often involve denaturing conditions during SF extraction from cocoons and employ processing techniques such as wet spinning,^8^ hand drawing,^9^ microfluidic systems,^10^ and electrospinning.^11^ However, these approaches often yield sub-optimal SF conformations that fail to form the native β-sheet crystalline structure, leading to inferior elastic properties compared to native SF fibers.^12–14^ To enhance β-sheet content and mechanical performance, post-processing of spun fibers in organic solvent baths based on ammonia,^15^ hexafluoroisopropanol/methanol,^16^ hexafluoroacetone hydrate/methanol,^17^ and formic acid/methanol^18^ have been used. However, these methods lead to the formation of highly compact β-sheets, resulting in either rigid and brittle, or very weak fibers that do not replicate the conformation and elasticity of native SF fibers.^13, 19–21^ Moreover, the use of such solvents is not compatible with biomedical applications due to potential toxicity for cells. Alternative approaches utilizing pH gradients, salt solutions (based on Na^+^, K^+^, Ca^2+^ and Cl^-^), and enzymatic crosslinking^22, 23^ have been explored to stabilize water-soluble SF without the use of organic solvents but still result in mechanically weaker materials compared to natural fibers.^24, 25^

To address these limitations, a novel processing method is proposed in this study to control the nano-to-microstructure of SF fibers. The method involves electrowriting of water-soluble SF solutions followed by post-processing in a sodium dihydrogen phosphate (NaH_2_PO_4_) solution (Figure 1). We hypothesized that the electrowriting method —a manufacturing process that uses electrical signals to control the flow of polymeric materials through an extrusion nozzle—can emulate the shear and elongation forces experienced in the spinning ducts of spiders and worms, thereby facilitating hierarchical self-assembly of SF. Additionally, the post-processing treatment in the phosphate-rich solution is expected to induce conformational transitions in the SF protein, specifically from random coil to α-helices and β-sheets, which could yield material properties different from those observed with more common techniques, such as methanol treatment.^19, 26^ Furthermore, SF electrowriting offers the potential to fabricate SF fiber constructs with controlled microgeometries, overcoming limitations of previous silk spinning methods in terms of patterning control, reproducibility, and consequent ability to mimic tissue morphology and biomechanics.^27, 28^ In this work, water-soluble SF inks with tuneable rheological properties were developed, and their compatibility for electrowriting of flat and tubular macrostructures with controlled microarchitectures was investigated. The effect of post-printing treatment with NaH_2_PO_4_ on restoring the secondary structure of SF protein was assessed, and the resulting mechanical properties of printed constructs were investigated under both quasi-static and dynamic conditions. Finally, the ability of SF fiber scaffolds to support the growth and viability of conditionally immortalized proximal tubule epithelial cells (ciPTEC) and conditionally immortalized glomerular endothelial cells (ciGEnC) was evaluated to assess the potential of SF fiber scaffolds for RM applications.

## 2. Results and discussion

### 2.1 SF biomaterial ink optimization for aqueous electrowriting

To achieve consistent and durable fibers for manufacturing ordered 3D structures using aqueous electrowriting, the development of inks with high viscosities is crucial.^37, 38^ Although increasing the concentration of SF beyond 20% resulted in inks with high viscosities, it led to electrical instabilities during jet formation due to increased electrical conductivity (Figure S1).^39^ To circumvent this issue, the SF concentration was fixed set at 20%, which yielded electrical conductivity values below 3 mS cm^-1^. Ink viscosity was further adjusted by varying the degumming time and incorporating a secondary polymer, poly(ethylene oxide) (PEO). Increasing PEO concentration up to 3% led to an increase in ink viscosity to a value of approximately 3 Pa s. The highest ink viscosity was observed when using the highest concentration of PEO (3%) in combination with the shortest SF degumming times (5 min). The measured viscosity values were consistent with those reported for other natural polymers used in solution-electrospinning processes.^40^ Among the tested ink formulations, the ink comprising a 20% SF solution, a short degumming time of 5 min, and 2.5 – 3% of PEO (SF5DT-3%) exhibited no electrical instabilities during jet formation, allowing for stable fiber deposition, and was therefore selected for further investigations (Figure 2B).

**Figure 2.**
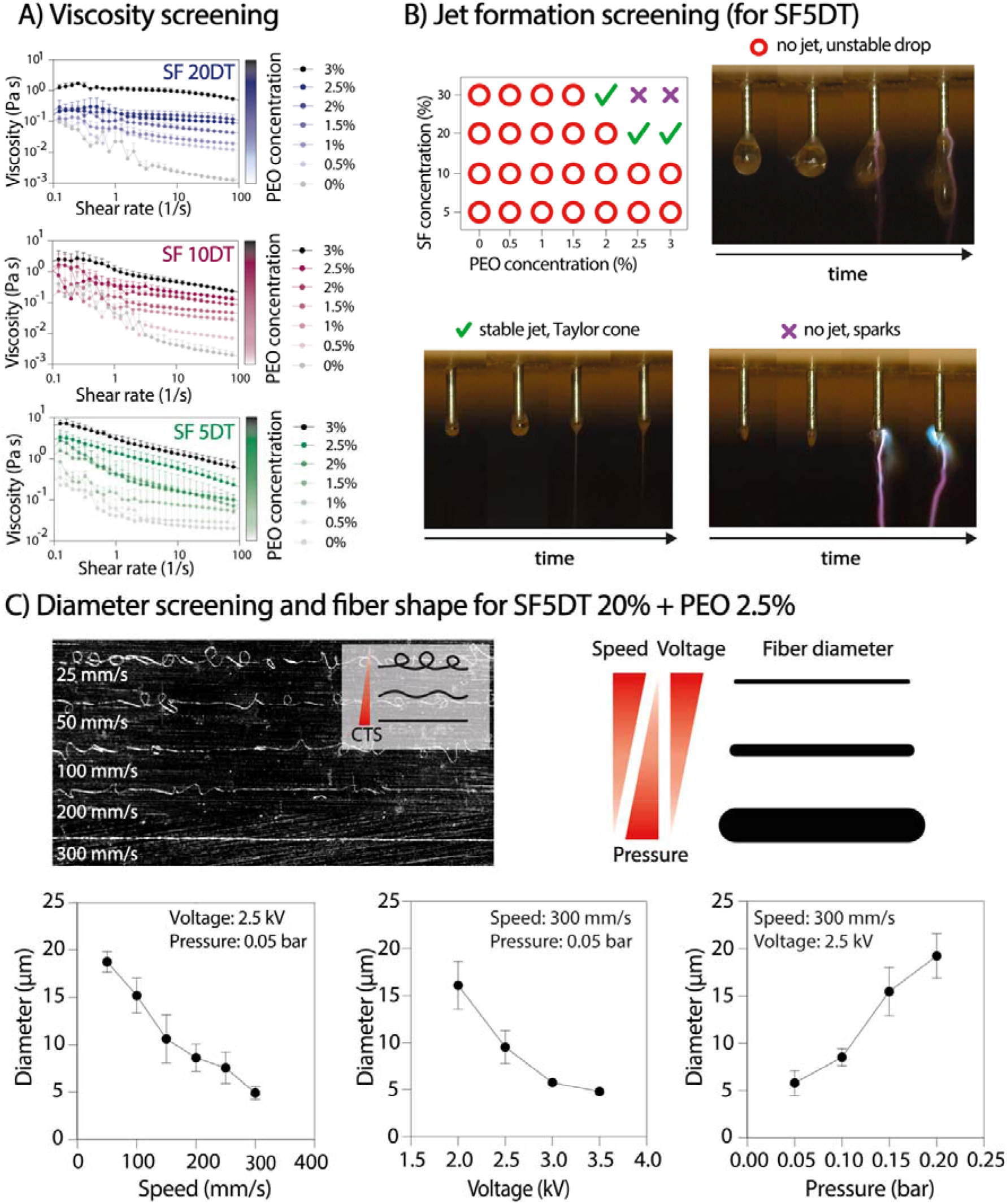
SF inks and electrowritten aqueous jet and fiber formation. A) Viscosity screening of SF solutions at 20% with increasing degumming time (5-, 10- and 20-min DT) and with increasing PEO concentration from 0 to 3% (n=3). B) Evaluation of SF ink on jet formation. C) Effect of key processing parameters of the aqueous electrowriting on fiber morphology and diameter (n=3).

Subsequently, the influence of key parameters in the electrowriting process on the morphology and diameter of SF fibers was investigated. By individually varying the collector speed, applied voltage, and pressure, straight fibers with diameters ranging from approximately 5 µm to 20 µm were obtained. A collector speed of 300 mm s^-1^ enabled the deposition of straight fibers (Figure 2C). Varying the collector distance (Cd) between 8 and 2 mm did not yield a significant effect on the fiber diameter, but a Cd value above 6 mm resulted in jet instabilities and poor control over fiber deposition. A combination of printing parameters, including a collector speed of 300 mm s^-1^, an applied voltage of approximately 2.5 kV, and pressure of 0.05 bar, was selected for subsequent tests due to its ability to generate a stable jet and produce straight fibers with a consistent diameter throughout the printing process.

### 2.2 Electrowriting of controlled microfiber shapes

To produce 3D scaffold structures with controlled microarchitectures, we evaluated the ability to stack SF microfibers for both flat and tubular shaped scaffolds (Figure 3). For flat scaffolds (Figure 3A-D), 400-μm and 1000-μm inter-fiber distances were evaluated, resulting in pore sizes with theoretical areas of 0.08 mm^2^ and 1 mm^2^, respectively. Optical and SEM imaging analysis confirmed the successful stacking of SF fibers up to 100 layers. As expected, a lower printing accuracy (as defined by quality number) was achieved for 100 layers scaffolds compared to those with 5 layers. Surprisingly, scaffolds with 400-μm inter-fiber distances resulted in higher quality number (= 0.78) than scaffolds with 1000 μm (∼ 0.62) (Figure 3B). Additionally, when the fiber spacing was set below 400 μm, unstable jets were observed due to repulsive forces generated between deposited fibers and polymer jet, which is consistent with previous findings.^41^ Consequently, the observed distances between the fibers were smaller than the theoretical values.^42^ For tubular scaffolds, winding angles of 60° and 90° were investigated, with theoretical pore areas of 1.82 mm^2^ and 3.16 mm^2^, respectively (Figure 3E-H).^43^ Optical and SEM analysis confirmed the accurate printing of tubular scaffolds with up to 100 stacked layers for both winding angles. Interestingly, printing accuracy did not lower with an increase in the number of stacked fibers, which is opposite to the observation for flat constructs (Figure 3F). Overall, the observed instabilities seemed to become more pronounced as the number of layers increased. Similarly to a previous study by Hai *et al*,^41^ we found that jet stability and the printing accuracy were also influenced by environmental factors such as temperature and humidity, which are challenging to control and seem to play a particularly bigger role during electrowriting of aqueous-based materials. Consequently, it may not always be possible to explain the differences in resolution between two theoretically identical scaffolds. Nevertheless, the small standard deviation in our printing accuracy data demonstrates that we can achieve good reproducibility.

**Figure 3.**
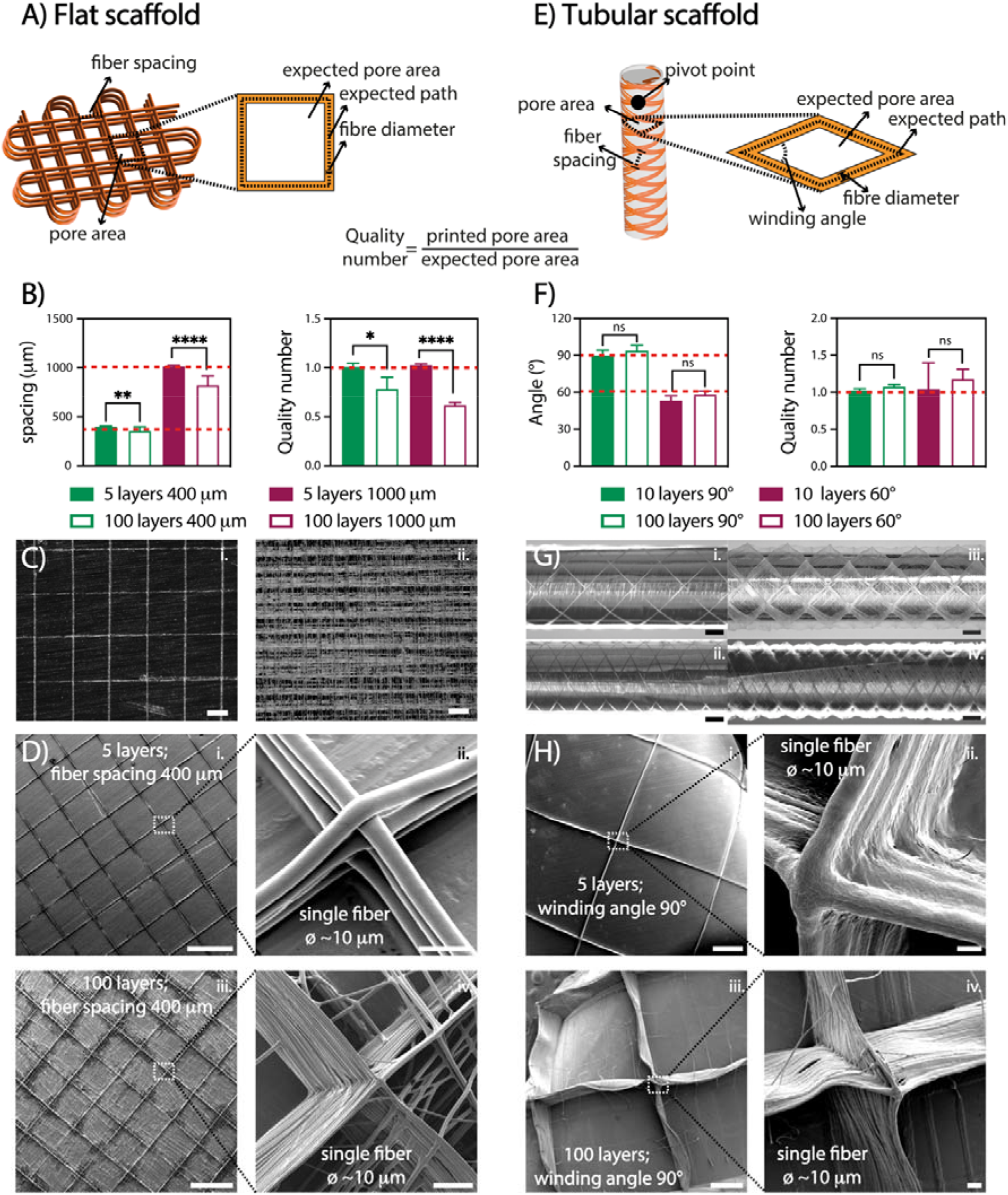
Electrowriting of controlled microfiber shapes. A) flat scaffold design and quality number, Q, definition. B) Inter-fiber distance spacing and quality number for flat scaffolds with 5 and 100 layers and with 400-μm and 1000-μm fiber spacing (n=10). C) Light microscope images of *i.* 5 and *ii.* 100 layers scaffold. Scale bars: 500 µm. D) SEM images of flat scaffolds; scale bars: *i.* and *iii.* 500 µm, *ii.* 10 µm, *iv.* 50 µm. E) Tubular scaffold design and quality number. F) Inter-fiber distance spacing and quality number for tubular scaffolds with 60° and 90° winding angle and with 10 and 100 layers (n=10). G) Light microscope images of tubular scaffolds with 90° (*i.* 5 layers and *iii.* 100 layers) and 60° (*ii.* 5 layers and *iv.* 100 layers) angle with 10 pivot points; scale bar: 1 cm. H) SEM images of tubular scaffolds; scale bars: *i.* and *iii.* 500 µm, *ii.* 10 µm, *iv.* 50 µm. (* = p < 0.05, ** = p < 0.01, **** = p < 0.0001).

### 2.3 Aqueous electrowriting of SF fibers and post-treatment restored conformation of **β**-sheet crystalline structure

The produced scaffolds exhibited a well-organized and precise 3D structure. However, the materials used were soluble in water, causing the printed scaffolds to lose their shape or dissolve within a few minutes upon immersion in aqueous solutions. To address this challenge, we explored the effect of elevated ion concentrations, taking inspiration from the natural crosslinking process of (spider) silk. Unfortunately, attempts with potassium, sodium, chloride, and sulfate ions resulted in unstable scaffolds that dissolved in water (data not shown). Notably, the treatment with NaH_2_PO_4_ led to physical crosslinking and the formation of insoluble SF fiber scaffolds over at least 28 days in PBS at 37°C (Figure S2). Therefore, the post-printing treatment of scaffolds in NaH_2_PO_4_ (SF-NaH_2_PO_4_) was compared to the standard methanol crosslinking method (SF-MeOH).^26^ Raman spectroscopy was used to investigate the secondary structure resulting from the different treatments (Figure 4A). Upon close examination of the spectra, it was observed that in un-crosslinked SF, the intensity of the β-sheet and α-helix bands was relatively low and similar to each other, indicating a random secondary structure of the SF protein.^23, 26^ However, both SF-MeOH and SF-NaH_2_PO_4_ exhibited a more intense, narrower β-sheet band (1200 - 1300 cm^-1^). In SF-MeOH, there was a shift in the amide I peak from 1666 cm^-1^ to 1662 cm^-1^, indicating an increase in β-sheet structures. In contrast, this shift was not observed in the NaH_2_PO_4_-treated sample.^26^ A comparison between SF-MeOH and SF-NaH_2_PO_4_ revealed that the β-sheet band in SF-MeOH was higher and narrower, while the α-helix band showed the opposite behaviour. These findings are consistent with previous studies suggesting that ionic charges can facilitate the alignment of silk molecules into β-sheets,^44^ and that phosphate ions in particular can stabilize protein helices, including those found in collagen, by enhancing inter- and intramolecular forces.^45^

**Figure 4.**
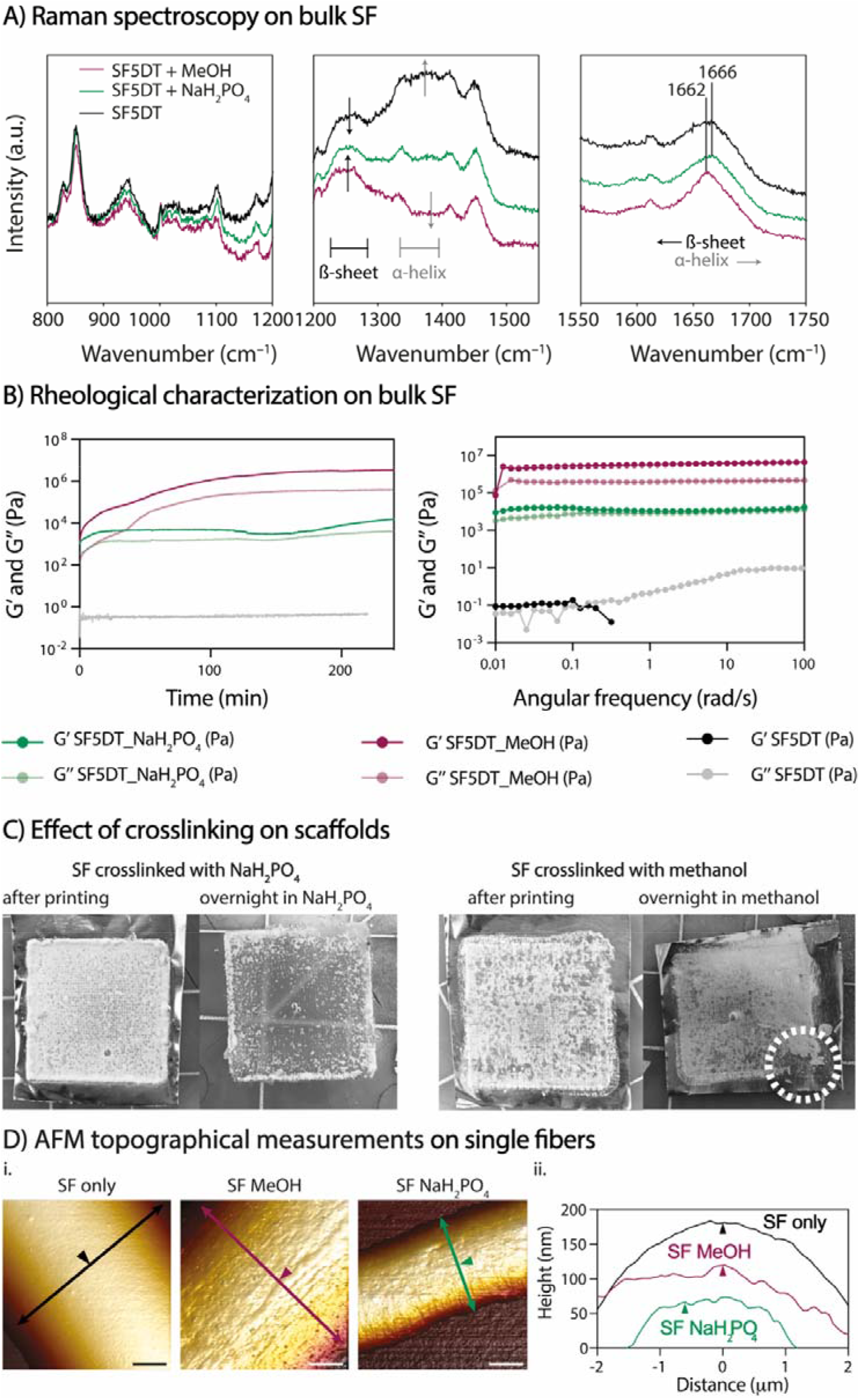
Crosslinking, β-sheet crystalline formation and topographical characterization. A) Raman spectrum of uncrosslinked SF, SF-MeOH, and SF-NaH_2_PO_4_. B) Rheological analysis of 20% SF + 2.5% PEO crosslinked with MeOH and with NaH_2_PO_4_, performed on bulk SF material. C) Effect of the different crosslinking treatment: stability of printed SF scaffolds in aqueous solutions. White dotted line indicates collapsed SF scaffold structure. D) Representative AFM images: *i.* 3D visualization of topography of single fibers and *ii.* height profiles of SF single fibers before (SF only) and after crosslinking treatment (MeOH and NaH_2_PO_4_); scale bars: *i.* 2 µm. Arrows are perpendicular to fiber axis and show location of height profile; arrowheads indicate equal positions in *i.* and *ii*.

The rheological properties of SF inks were also investigated for both crosslinking methods, and time sweep measurements demonstrated that crosslinking occurred almost instantaneously for both methods. The shear modulus of SF-NaH_2_PO_4_ ink was lower (almost two orders of magnitude) than for SF-MeOH, which could be attributed to a slightly higher β-sheet content in SF-MeOH (Figure 4B). Degumming time was also found to affect the moduli of individual SF inks when exposed to both crosslinking methods. As expected, an increase in degumming time led to a decrease in SF-ink modulus (Figure S3). While both bulk SF-MeOH and SF-NaH_2_PO_4_ crosslinked gels are stable in PBS over time (Figure S4), printed SF-MeOH scaffolds exhibited high brittleness, which hindered their handling. Upon comparing scaffolds before and after crosslinking with NaH_2_PO_4_ and MeOH (Figure 4C), it was observed that upon immediate contact with NaH_2_PO_4_ solution, the scaffolds turned opaque, and they eventually detached from the aluminum surface after soaking overnight. In contrast, scaffolds immersed in MeOH were brittle to the touch and impossible to remove from the aluminum surface without breaking them. The ease of handling of SF-NaH_2_PO_4_ scaffolds can be attributed to a less regular sequence of β-sheet structures, which allows for potentially high fiber elongation, as observed in native spider silk.^46^

To assess the topography of individual SF fibers exposed to different crosslinking conditions, AFM analysis was performed (Figure 4D). The surface texture of crosslinked SF fibers exhibited nano-striations that aligned with the fiber’s main axis (Figure 4D and Figure S5). These striations likely result from shear-induced chain alignment during the aqueous electrowriting process, becoming visible on the surface after crosslinking due to the evolution of SF secondary structure. Fiber dehydration upon exposure to either methanol or a phosphate-rich solution may contribute to the formation of multiple nanometer and micrometer-scale fibers, which that were not visible in the un-crosslinked SF. Additionally, the 3D microstructure of SF-NaH_2_PO_4_ tubular scaffolds was analyzed by micro-computed tomography (µCT; Figure S6), revealing a mean strut thickness of 166 ± 48 µm and a scaffold porosity of 97.1%.

### 2.4 Mechanical analysis

Uniaxial tensile tests were conducted on SF-NaH_2_PO_4_ flat scaffolds with fiber spacing of 400 μm and 1 mm, as well as on tubular scaffold with 90° and 60° winding angle (both 100 layers) to investigate the impact of crosslinking on SF scaffold elasticity. After NaH_2_PO_4_ treatment, both flat and tubular scaffolds were tested along the main plane and radially, respectively, under uniaxial tension while immersed in PBS to maintain equilibrium swelling of crosslinked SF (Figure 5A and 5B). The results showed that the stiffness of the flat scaffold with 400-µm spacing was significantly higher (233.5 mN) than that of the 1-mm spacing (64.95 mN) (Figure 5Aiii). The ratio of stiffness between 400-µm and 1000-µm scaffolds was greater than the ratio of number of cross-sectional fibers. This suggests that stiffness is not solely dependent on the number of fibers in the cross-sectional area, but also on the reinforcing effect between the SF molecule chains. This effect increased with the number of printed fibers, particularly for the 400-µm inter-fiber spacing group. Tubular scaffolds exhibited a toe region at low strains due to gradual pore elongation along the tensile force. The 90° winding angle scaffolds showed a wider toe region but lower stiffness (145.6 mN) compared to the 60° winding angle scaffolds (818.7 mN), which had fewer elongated pores.

**Figure 5.**
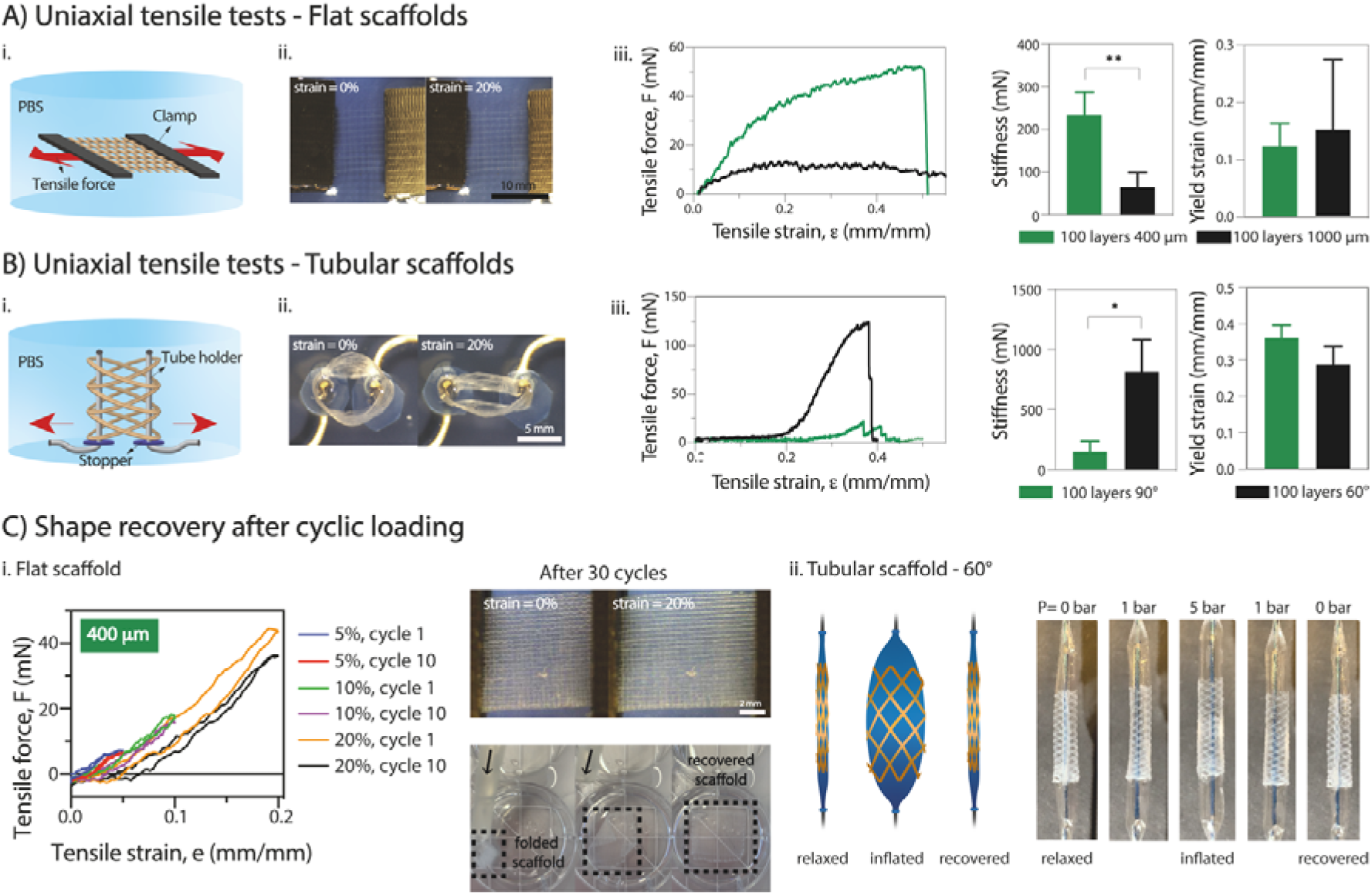
Mechanical characterization. A) *i.* Side-view schematics and *ii.* top-view photographs of the uniaxial tensile testing setup for flat scaffolds; *iii.* Representative uniaxial tensile curves and tensile properties of flat scaffolds (n = 3–5; unpaired t-test with Welch’s correction). B) *i.* Side-view schematics and ii. top-view photographs of the uniaxial tensile testing setup for tubular scaffolds; *iii*. Representative uniaxial tensile curves and *ii.* tensile properties of tubular scaffolds (n = 3–5; unpaired t-test with Welch’s correction). C) Shape recovery properties of *i.* flat and *ii.* tubular scaffolds; images obtained while applying unidirectional cyclic stimuli, and tests with catheters for *i.* flat scaffold (scale bar: 1 cm) and *ii.* tubular scaffold (scale bar: 400 µm).

Invasive surgical procedures, as demonstrated in previous studies, are currently utilized for the application of cardiac patches,^30^ vascular stents,^48^ and kidney stents^49^. However, these procedures are not ideal, and less invasive approaches are preferred. To showcase the versatility of our scaffolds, we tested their shape recovery properties, both for flat and tubular shapes. Two tests were performed: cyclic strain-recovery tests on the scaffold and a catheter analysis to simulate the mechanical environment to which the scaffold may be exposed during microsurgery (Figure 5C). Uniaxial cyclic tests were performed on flat scaffolds (Figure 5Ci), involving 10 cycles of constant strain-recovery with deformations up to 20%. The scaffold showed complete elastic behavior for deformations up to 10%, recovering its original shape between cycles without plastic deformation. However, at deformations of 20%, an onset of plastic deformation was observed, resulting in incomplete shape recovery (Figure 5Ci and Figure S7i). Notably, the SF-NaH_2_PO_4_ scaffold exhibited these shape recovery properties, while the SF-MeOH scaffold did not, as it appeared to be more fragile and prone to rupture. Similar behavior was observed for tubular scaffolds, although the uniaxial cyclic tests did not fully replicate the physiological conditions due to the absence of radial deformation, as typically seen in blood vessels and kidney tubules. Flat scaffolds with 400-µm and 1000-µm inter-fiber spacing were found to be optimal for flowing inside a tube catheter (Ø = 500 µm), showing no signs of rupture, kink formation, or plastic deformation (Figure 5Ci). Furthermore, tubular scaffolds were subjected to a balloon catheter test to assess radial shape recovery (Figure 5Cii). The deflated balloon was introduced into the tube and then inflated to full extension (pressure up to 5 bar). Catheter swelling caused the scaffold fibers to stretch radially without rupture, and subsequent deflation of the catheter enabled the scaffold to restore its original shape and size. The swelling/deswelling of the balloon was applied multiple times (3 times) without any observation of plastic deformation (initial tube diameter = 3 mm; tube diameter after 3 cycles ∼ 3 mm) (Figure S8).

### 2.5 Flat and tubular scaffolds support cell growth

To demonstrate the suitability of SF scaffolds crosslinked with NaH_2_PO_4_ for *in vitro* cell culture, we conducted stability tests over a period of 28 days at 37°C in PBS. Notably, no visible degradation was observed (Figure S2). Subsequently, to investigate whether SF fiber scaffolds could support cell growth, proximal tubular (ciPTEC) and glomerular endothelial (ciGEnC) cells were seeded on both flat and tubular scaffolds. While both cell types were not able to attach to the SF fiber scaffolds without a biofunctionalization step (Figure S9), L-DOPA coating enabled cell attachment and monolayer formation (Figure 6). Both ciGEnC and ciPTEC formed monolayers on the flat and tubular scaffolds within 7-14 days after initial seeding. Furthermore, ciPTEC and ciGEnC grown on flat scaffolds showed high viability (>95%), deposition of type-IV collagen and limited production of type-I collagen. F-actin directionality indicated preferential cell alignment along scaffold microstructure, observed with a 90° orientation (Figure 6). Similarly, ciPTEC and ciGEnC grown in tubular scaffolds formed aligned monolayers and deposited type-IV collagen and a limited amount of type-I collagen, indicating that cells began depositing their own extracellular matrix. Furthermore, the presence of cell-specific markers α-tubulin (ciPTEC) and CD31 (ciGEnC) confirmed cell proliferation within the scaffolds (Figure S10).

**Figure 6.**
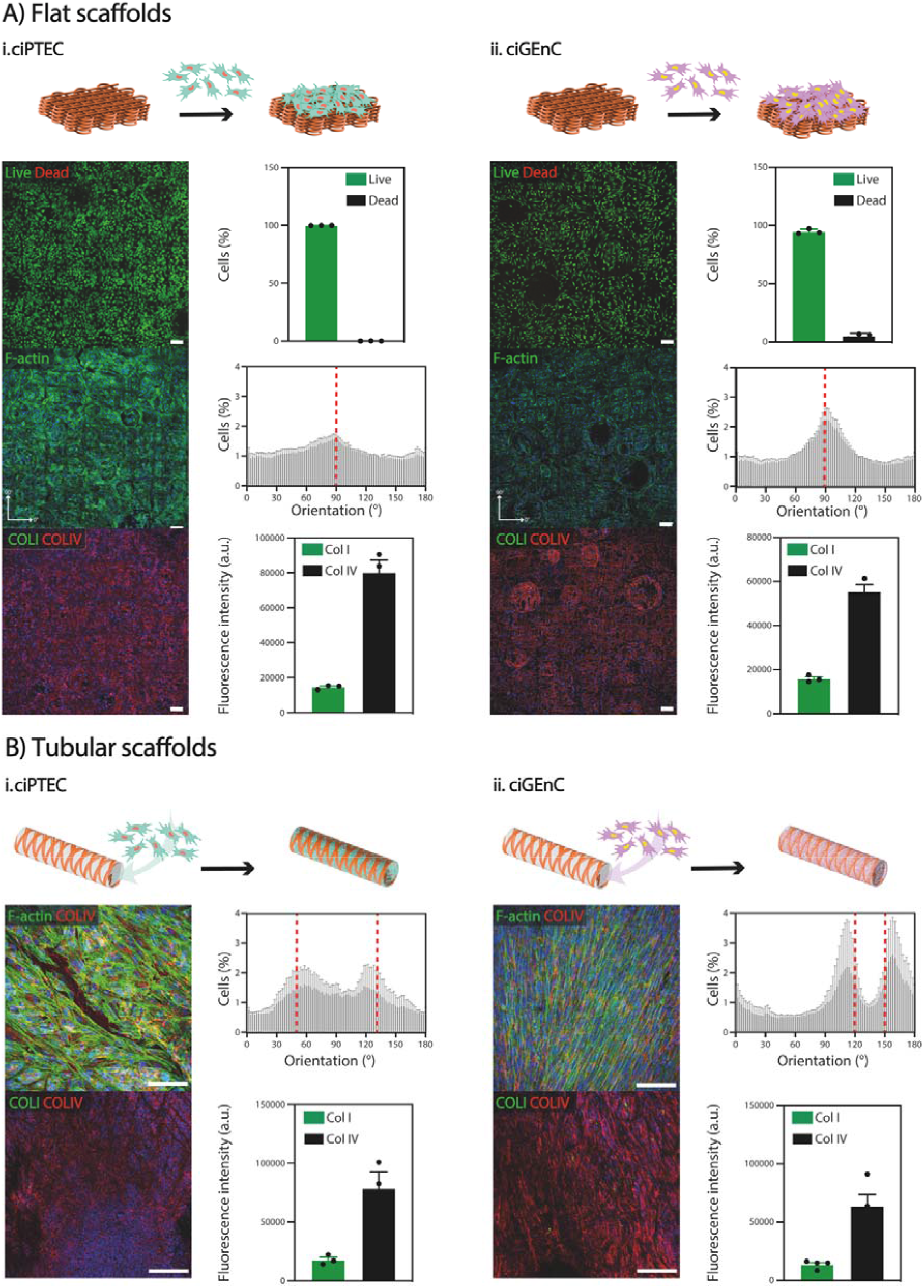
Cell viability and growth in A) flat (400 μm, 100 layers) and B) tubular (60°, 100 layers) scaffolds for *i.* ciPTEC and *ii.* ciGEnC. Scale bar: 100 μm.

Our findings are consistent with existent literature. Previous studies have explored the influence of various scaffold topographical features, such as scaffold geometry, pore area and pore geometry, on the behaviour of kidney cells.^50^ For instance, Van Genderen *et al.* demonstrated that the geometry of polycaprolactone-based fiber scaffolds influenced cell alignment, thereby impacting expression of genes with key roles in transport functionality.^51^ Furthermore, they observed that the maximal scaffold pore size possible to ensure complete pore filling was cell type-dependent, with ciPTECs showing the ability to spread widely and with high adherence over porous scaffolds.

## 3. Conclusions

In this study, an aqueous-based electrowriting technique was developed to create microfiber scaffolds made from silk fibroin with distinct mechanical properties. The resulting flat and tubular scaffolds exhibited microfiber sizes with diameters as small as 5 µm, along with well-defined square-based and crosshatch microarchitectures. A crosslinking method based on a aqueous solution of sodium dihydrogen phosphate was also introduced, which partially restored the β-sheet structure of silk fibroin, resulting in increased elasticity compared to conventional crosslinking methods involving organic solvents. The fiber scaffolds showed unique shape recovery properties and stability in aqueous environments. Moreover, when coated with L-DOPA, the scaffolds supported the growth of ciPTEC and ciGEnC cells, which formed uniform and aligned monolayers and produced type-IV collagen, indicating healthy epithelial tissue. This innovative fiber-based processing approach holds great promise for creating silk fiber scaffolds with unique elastic properties, making them highly suitable for regenerative medicine applications.

## 4. Experimental Section

### Materials

SF was extracted from *Bombix mori* cocoons (kindly provided by Evrosilk; Czech Republic) as described elsewhere.^29^ Briefly, the cocoons were degummed at 100°C in a 0.02 M sodium carbonate (Na_2_CO_3_; Sigma Aldrich) solution for 5, 10 or 20 minutes, dissolved in a 9.3 M lithium bromide (LiBr; Acros Organics) solution for 4 h at 60°C and then dialyzed at 4°C against pure water for 4 days using cellulose dialysis tubes (MWCO 3.5 kDa, Sigma Aldrich). Subsequently, the diluted solution was concentrated with inverse dialysis at 4°C using cellulose dialysis tubes (MWCO 3.5 kDa, Sigma Aldrich) against a 40% PEG (6 kDa; Sigma Aldrich) aqueous solution for 16 h to obtain a SF ink concentration between 15 and 20%. Three ink solutions were studied SF5DT, SF10DT and SF20DT, where the number indicates the degumming time (minutes of boiling). Poly(ethylene oxide) (PEO) (10 kDa; Sigma Aldrich) was added prior to printing, in quantities ranging from 0.5 to 3%.

### Ink material characterization

The viscosity of 20% SF aqueous solutions with increasing amount of PEO (10 kDa; Sigma Aldrich), was analysed with a rheometer (Discovery HR2, TA Instruments, New Castle, DE, USA) using a cone-plate geometry (20 mm – 1°) performing a flow sweep at 25°C with shear rate from 0.1 to 1000 s^-1^. The electrical conductivity of the solution was measured with an electroconductive-meter (Consort C861). Distilled water was used as a standard and calibrated to 1413 μS/cm. The temperature was set to 25°C and automatic temperature compensation was applied. The electrodes were washed with DI water before every measurement and afterwards immersed in the SF solution for 1 min (n = 3).

### Aqueous electrowriting of SF fibers and fiber scaffolds

Electrowriting was performed using an in-house built set up that allows for both flat and tubular scaffolds manufacturing as described before.^30, 31^ SF5DT aqueous solution (20%) with 2.5% PEO ink was poured in a 3 ml glass syringe (Fortuna optima Ganzglasspritze, Poulten & Graf GmbH) with a 27G metal needle (Unimed), connected to a sealed hose delivering pressurized nitrogen (VPPE-3-1-1/8-2-010-E1, Festo). The SF solution was electrified using a high voltage source (Heinzinger, LNC 10000-2neg) and collected either onto a grounded collector plate (x-y) or onto a rotating aluminium mandrel (Ø = 4 mm and 3 mm) mounted on a x-y axis, both covered in aluminium foil and controlled by an advanced motion controller Motion Perfect v5.0.2 (Trio Motion Technology Ltd.). The polymer solution processing compatibility was systematically investigated, specifically by tuning the voltage (V), the pressure (p), and the collection distance (Cd). The collector speed (Scol) was studied differently for flat and tubular scaffolds as previously described.^32^ For both flat and tubular scaffolds, fiber diameter and morphology were investigated by changing the mentioned parameters between the following values: V = [1.5 - 6] kV, p = [0 – 0.1] bar, Scol = [20 - 400] mm/s, and Cd = [2 - 6] mm, one parameter at a time. The optimized SF flat scaffolds were manufactured using the following parameters: V = 2.5 kV, p = 0.05 bar, Scol = 300 mm s^−1^, and Cd = 6 mm. Organized scaffold meshes (40 cm^2^) with squared microstructures (fibers spacing 400 μm and 1 mm) were fabricated. The SF tubular scaffolds were manufactured using the following parameters: V = 4.8-5.3 kV; p = 0.1 bar; Scol = 100 mm s^−1^ and Cd = 6 mm. Tubular scaffolds with winding angles 60° and 90° and corresponding to a square or rhomboid microarchitecture with pore sizes of 1.82 mm^2^ and 3.16 mm^2^ respectively, were fabricated.

### Electrowritten constructs imaging before post processing

Printed SF scaffolds were characterized first with a stereo microscope (Olympus SZX7) and then with Scanning Electron Microscope (SEM, Quanta3D FEG, Thermofisher Scientific, Eindhoven, The Netherlands). For SEM, all scaffolds were coated with Pt (30s, 30 mA) using a Q150T S high resolution sputter coater (Quorum) and imaged at 5 kV. All the images were analysed with Fiji ImageJ software. The number of fibers used for the diameter measurements was at least n = 10. Fiber spacing, winding angle and quality number were used as references to characterize the accuracy of printed scaffolds. The quality number is identified as the ratio between the experimental and theoretical pore area; in an ideal condition, these two values are identical, and the quality number is 1.

### Post processing treatment

SF scaffolds were immersed in either a solution of methanol (Sigma Aldrich) or 2 M aqueous solution of sodium dihydrogen phosphate (NaH_2_PO_4_; Sigma Aldrich) overnight, after which the scaffolds were washed with 10 mL of DI water 5 times and then kept in DI water.

### Material properties after crosslinking, β-sheet formation and electrowritten fibers topography

To assess SF crosslinked material properties, rheological characterization of SF5DT 20% - PEO 2.5% solution with either methanol or 2M NaH_2_PO_4_, was performed with a rheometer (Discovery HR2, TA Instruments, New Castle, DE, USA) fitted with a plate-plate geometry (20 mm) with 300 μm gap. A time sweep with angular frequency 10 rad/s and strain 1% was conducted for both types of samples at 25°C. Both storage modulus (G′) and viscous modulus (G″) were recorded as function of time. The G’ and G’’ were also evaluated as a function of the strain (oscillation amplitude, T = 25°C, angular frequency = 10 rad s^-1^, strain from 0.1 to 100%) and as a function of the angular frequency (oscillation frequency, T = 25°C, strain = 1%, angular frequency from 0.1 to 100 rad s^-1^). β-sheet formation was evaluated with Raman spectroscopy using a Renishaw InVia Raman microscope with a 785 nm laser. The spectra were collected between 100 - 3200 cm^−1^ using a relatively low laser power (0.23 W) to avoid light-induced sample damage with an integration time of 5 minutes and 30 accumulations per spectrum (total acquisition time of 2.5 hours). The topography of single printed fibers was assessed by atomic force microscopy (AFM). A Multimode Nanoscope IIIa (Veeco) was used to record AFM height and peak force error images of SF fibers deposited on glass coverslips. Single fibers were immersed in either methanol or 2M NaH_2_PO_4_ for 12 h and air-dried before imaging. AFM scans were collected in the ScanAsyst large-amplitude mode at 23°C in air with silicon tips on nitride levers (ScanAsyst-Air, Veeco, 50–90 kHz, 0,4 N/m), controlled using Nanoscope 8 (Bruker), and processed using NanoScope Analysis 1.40 (Bruker).

### Micro-computed tomography (µCT)

NaH_2_PO_4_-crosslinked SF tubular scaffolds (150 layers, 60° winding angle) were immersed in a 1% w/v solution of phosphotungstic acid (Sigma Aldrich) as contrast agent for 3 h. The scaffolds were then thoroughly rinsed and imaged in PBS. μCT images were taken on a μCT100 system (Scanco Medical) with the following scanning parameters: voxel size = 36.8 μm, energy level = 70 kV, intensity = 114 μA, and integration time = 200 ms, with no filter, over a region of interest (ROI) of 162 slices. Noise reduction was performed with a constrained Gaussian filter, with a support value of 1.0 and a sigma width of 0.8 voxel. After segmentation, Image Processing Language (IPLFE v2.03, Scanco Medical) was used to analyse the 3D reconstructed models. Strut thickness was estimated as trabecular thickness (Tb.Th) by the distance transformation method,^33^ while porosity was calculated as P = 1 – (material volume excluding porosity / total volume including porosity).^34^

### Mechanical characterization

Uniaxial tensile tests were performed at 23°C in PBS using a BioTester 5000 device (CellScale) with a 1.5 N load cell. Flat scaffolds (width: 10 mm, active length: 10 mm) were tested monotonically at a strain rate of 10% min^-1^ (n ≥ 3); or under cyclical conditions at a loading-unloading frequency of 0.1 Hz with 10 cycles at 5% max. strain, followed by 10 cycles at 10% max. strain, and 10 cycles at 20% max. strain (n ≥ 3). Tubular scaffolds (axis length: 10 mm, inner diameter: 4 mm) were mounted on custom wire holders to allow for uniform radial deformation and tested monotonically at a strain rate of 10% min^-1^ (n ≥ 3). Force-displacement curves were recorded using LabJoy (CellScale). Force data were processed with a moving average filter (size: 15 datapoints). Stiffness constant (with units of force) was calculated from least square fitting of the linear region slope in force-strain curves. Yield strain was set as the upper value of the range used for stiffness calculations. Decrease in peak force was quantified as: 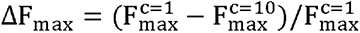, where 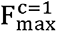 and 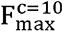 are the maximal forces during cycle 1 and 10, respectively, of each strain step. For delivery tests, a flat SF scaffold (30x30 mm, 100 layers) was passed through a catheter (Ø inner = 500 μm) with a constant flow of PBS. The scaffolds were collected in a petri dish and imaged. For the balloon catheter test, a tubular scaffold (tube length: 17.41 mm, Ø = 3 mm, winding angle: 60°, 100 layers) was introduced into a balloon catheter (XTRM Way 3, Blue Medical), with a diameter at maximum pressure of 5 mm. The deflated catheter was inserted into the tubular scaffold, and pressure was gradually increase up to 5 bar. At pressure intervals of 1 bar, images of the scaffold were taken to assess its diameter.

### Cell culture

Human conditionally immortalized proximal tubular epithelial cells (CiPTEC - obtained via Cell4Pharma, Nijmegen, The Netherlands) and glomerular endothelial cells ciGEnC were cultured as previously described.^35, 36^ In short, both cell lines were cultured in T75 flasks (Greiner Bio-One, Alphen aan den Rijn, The Netherlands) in their respective culture medium. CiPTEC were cultured in Dulbecco’s modified eagle medium/HAM’s F12 without phenol red (Thermofisher Scientific, Paisley, UK), with added insulin, transferrin, selenium (all 5 μg/ml), hydrocortisone (35 ng/ml), epidermal growth factor (10 ng/ml), tri-iodothyronine (40 pg/ml) (Sigma-Aldrich, Saint Louis, MO, USA), 10% fetal bovine serum (v/v, FBS, Greiner Bio-One), and 1% penicillin/streptomycin (v/v, Gibco, Thermofisher Scientific) to prevent infections. CiGEnC were cultured in Endothelial Cell Basal Medium-2 (Lonza) containing Microvascular Endothelial Cell Growth Medium-2 SingleQuots Kit (EGM-2 MV, Lonza). Both cell lines proliferate at the permissive temperature of 33°C and maturate at 37°C.

### Cell seeding

Tubular scaffolds were sterilized in a laminar flow cabinet by washing 3 times for 15 minutes in 5% penicillin/streptomycin (v/v, Gibco, Thermofisher Scientific) in phosphate buffered saline (PBS, Lonza). Thereafter, scaffolds were sterilized by UV exposure in a laminar flow cabinet (365 nm for 30 min). After sterilization, scaffolds were functionalized using 2 mg ml^-1^ L-3,4-dihydroxyphenylalanine (L-DOPA, Sigma-Aldrich) dissolved in 10 mM tris(hydroxyethyl)aminomethane (Tris) pH 8.5 buffer (Sigma Aldrich). After dissolving L-DOPA, it was put at 37°C for 45 minutes. After 45 minutes L-DOPA was filter-sterilized and used for functionalization. Scaffolds were immersed in L-DOPA solution at 37°C for a minimum of 4 hours. Tubular scaffolds were washed 3 times using PBS after the coating procedure. CiPTEC or ciGEnC were seeded in the tubular scaffolds (tube length: 17.41 mm, Ø = 3 mm, winding angle: 60°, 100 layers) at concentrations 15 × 10^6^ cells ml^-1^ and 30 × 10^6^ cells ml^-1^ respectively, using a positive displacement pipette. Two hours after seeding, scaffolds were turned 180°. Four hours after seeding, the respective culture medium was added. Scaffolds were then cultured until confluency at 33°C, followed by 7 days of differentiation at 37°C. Culture medium was refreshed every 2-3 days.

### Cell analysis and immunocytochemistry

Live/dead staining was performed by rinsing cells with PBS, followed by incubation with 2 μM calcein-AM and 4 μM ethidium homodimer-1 (Invitrogen) for 30 min to examine viability. For immunocytochemistry, cells were fixed for 10 min with 4 wt% paraformaldehyde (Pierce, Thermofisher Scientific) in PBS. Afterwards, cells were permeabilized using 0.3 wt% triton X-100 (Sigma-Aldrich) in PBS for 30 min and exposed to blocking buffer (2 wt% FBS, 0.5 wt% bovine serum albumin (Sigma Aldrich) and 0.1 wt% Tween-20 (Sigma Aldrich) in PBS) for 60 min at RT. Cells were incubated with primary and secondary antibodies diluted in blocking buffer for 1.5 hour and 1 hour at RT respectively. Table S1 contains a list of primary and secondary antibodies used. Confocal Leica TCS SP8 X microscope and software Leica Application Suite X (Leica) was used to examine immunofluorescence. Images were analysed using ImageJ (National Institutes of Health, USA).

### Statistical analysis

All data are shown as mean ± SD, unless otherwise stated. Statistical significance was tested by unpaired t-test with Welch’s correction, or two-way ANOVA with Tukey’s multiple comparisons test, as noted in each case. All statistical analysis was performed with Prism 9 software (GraphPad).

## Supporting Information

Provided in the file attached.

## Acknowledgements

The authors would like to thank Jim de Ruiter from the department of Inorganic Chemistry and Catalysis (Faculty of Science, Utrecht University, Utrecht, the Netherlands) for performing the Raman spectroscopy measurements. They also would like to thank the group of Prof. J. van der Vlag (RadboudUMC, Nijmegen, The Netherlands) and Prof. S. Satchell (Bristol Medical School (THS), Bristol, UK) for providing glomerular endothelial cells (ciGEnC). The authors gratefully thank the following agencies for their financial support: the European Union’s Horizon 2020 research and innovation programme, the Gravitation Program “Materials Driven Regeneration” (024.003.013), the Marie Skłodowska-Curie Actions (RESCUE #801540), the strategic alliance Utrecht Medical Centre Utrecht – Technical University Eindhoven and the Reprint project (OCENW.XS5.161) by the Netherlands Organization for Scientific Research.

