## Supplemental Information for "Microstructured silk-fiber scaffolds with enhanced stretchability"

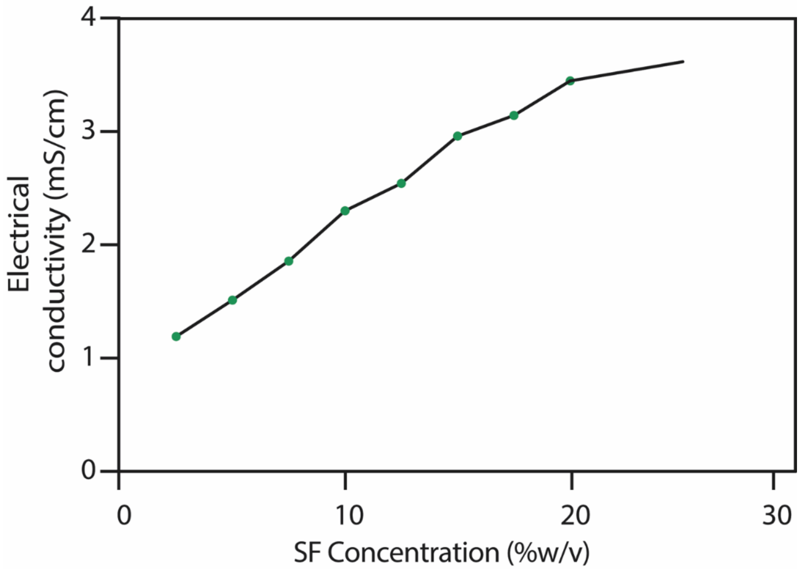


**Figure S1.** Electrical conductivity of SF5DT aqueous solutions at different concentrations.

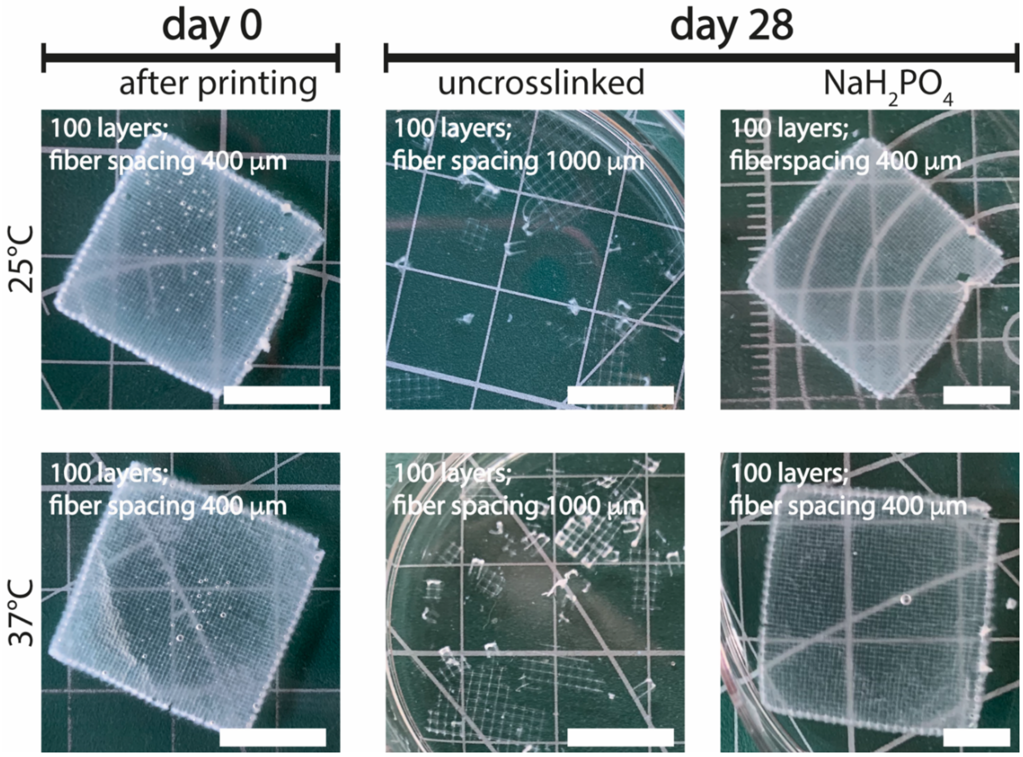


**Figure S2.** Stability of flat scaffolds at 25°C and 37°C in PBS over 28 days with and without $NaH_2PO_4$ treatment; scale bars: 1 cm.

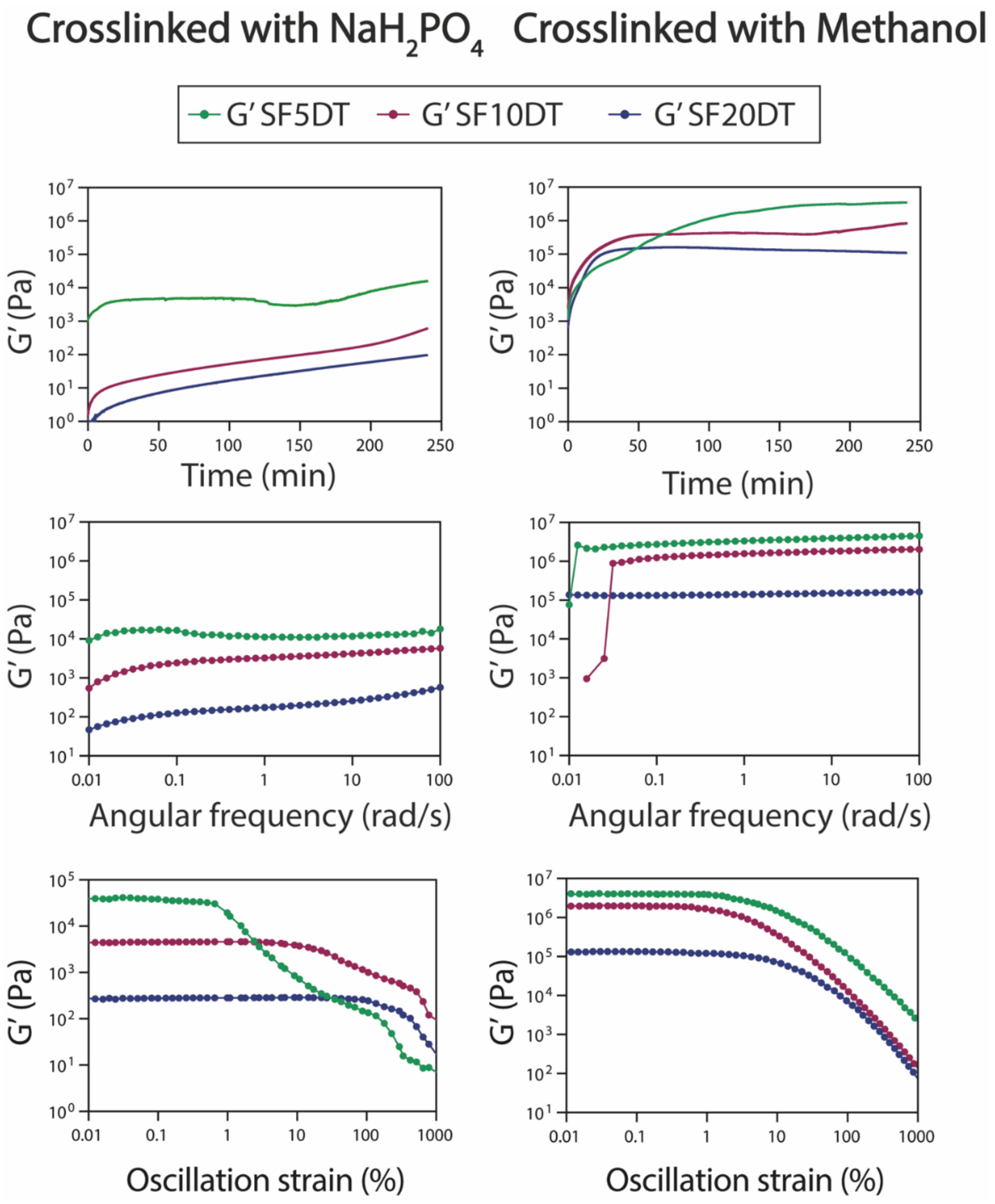


**Figure S3.** Rheological characterization of SF (at 20% w/v) with different degumming times, crosslinked with sodium dihydrogen phosphate ($NaH_2PO_4$; left panel) and methanol (MeOH; right panel).

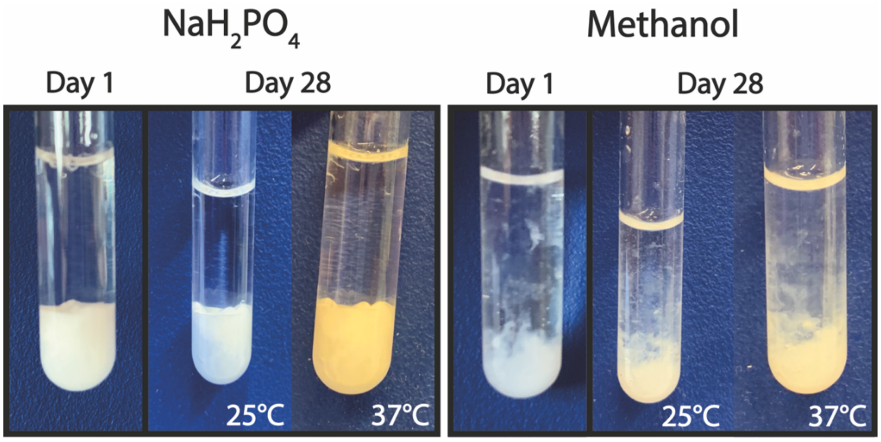


**Figure S4.** Stability of bulk gels crosslinked with $NaH_2PO_4$ and methanol over 28 days at 25°C and 37 °C.

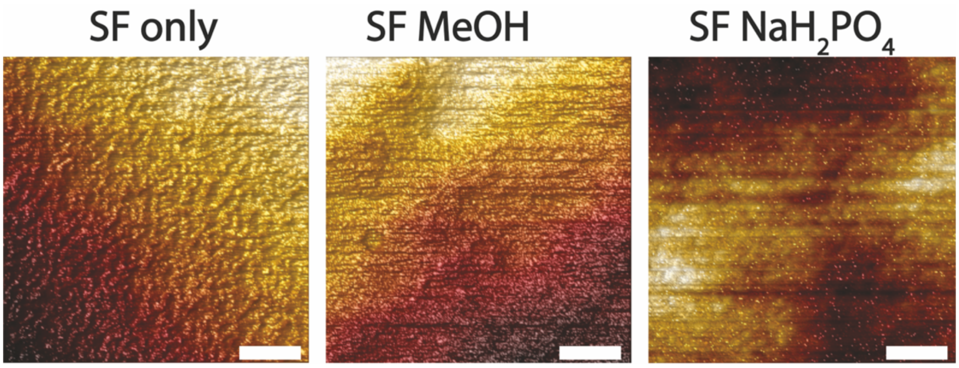


**Figure S5.** AFM 3D visualization of topography of single SF fibers before crosslinking (SF only) and after crosslinking (SF MeOH and SF $NaH_2PO_4$); scale bars: 200 nm.

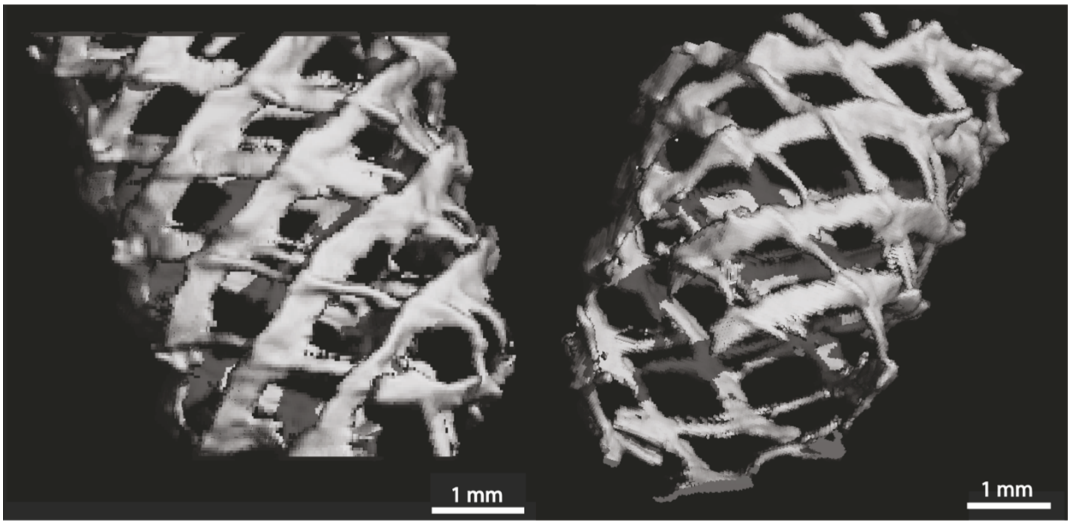


**Figure S6.** μCT reconstruction of SF-$NaH_2PO_4$ tubular scaffold (after printing, with post-processing treatment).

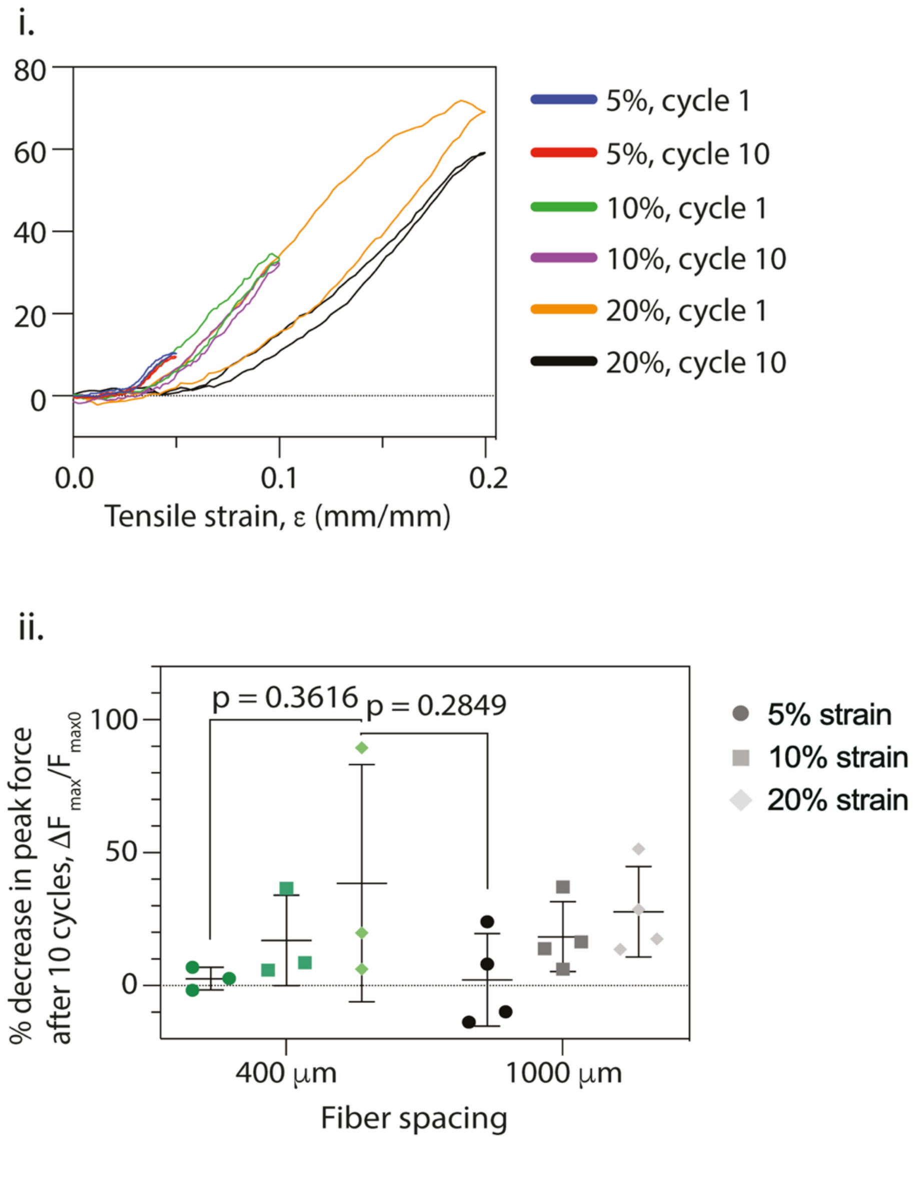


**Figure S7.** Cyclic tensile testing of flat scaffolds with inter-fiber spacing of 1000 μm. i) Representative curves, and ii) decrease in maximum tensile force of scaffolds after 10 cycles with 5% maximum strain (cycles 1–10), followed by 10 cycles with 10% maximum strain (cycles 11–20), and 10 cycles with 20% maximum strain (cycles 21–30) (n = 3–5; two-way ANOVA with Tukey's multiple comparisons test).

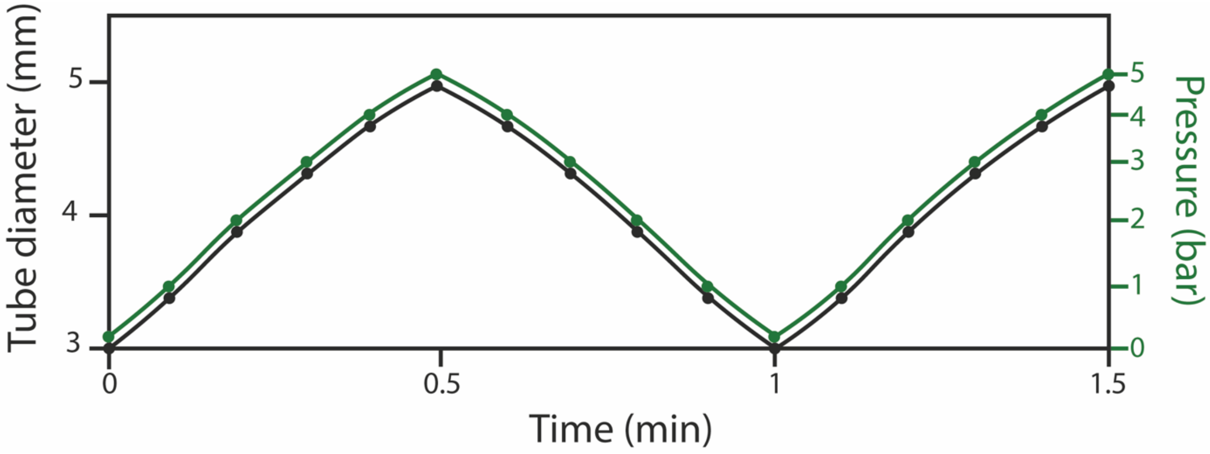


**Figure S8.** Variation of tubular scaffold diameter with the swelling and deswelling of a balloon catheter.

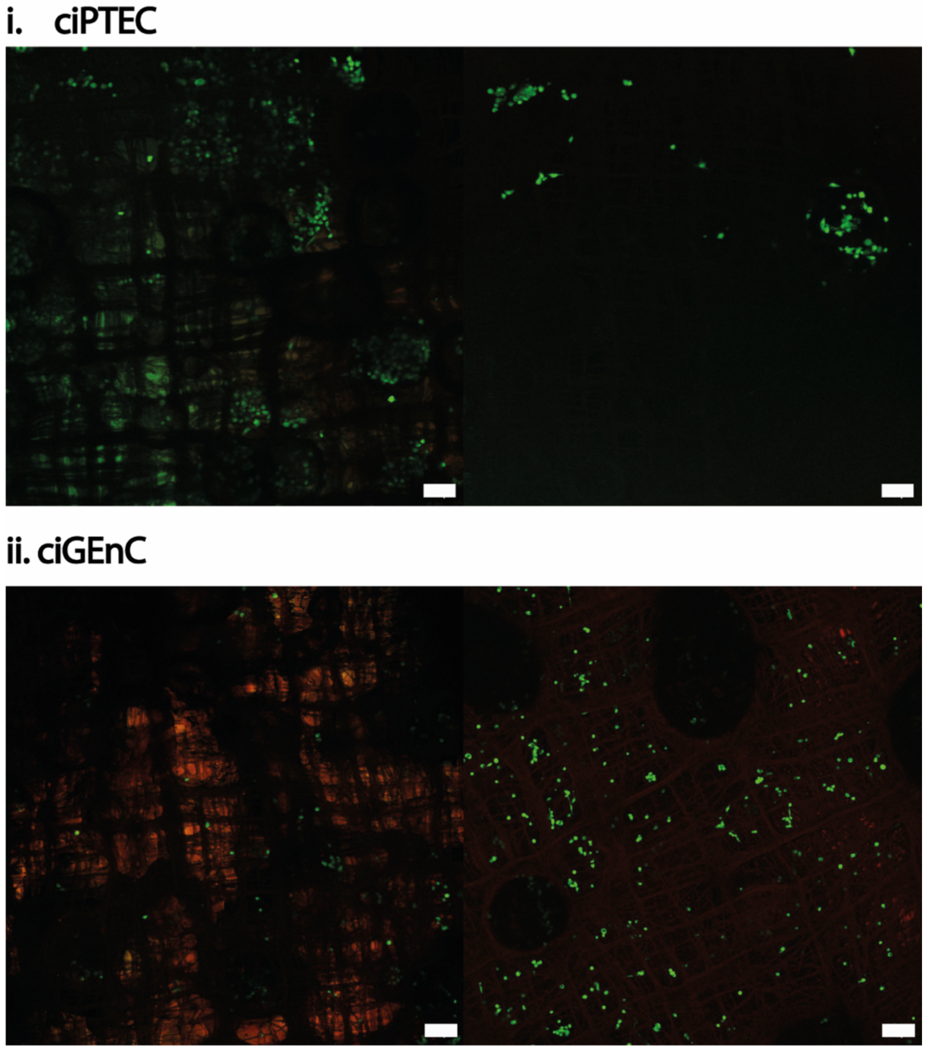


**Figure S9.** Human conditionally immortalized proximal tubular epithelial cells (ciPTEC) and glomerular endothelial cells (ciGEnC) cultured on non-coated flat scaffolds and stained for live (green) and dead (red) cells; scale bars: 100 μm.

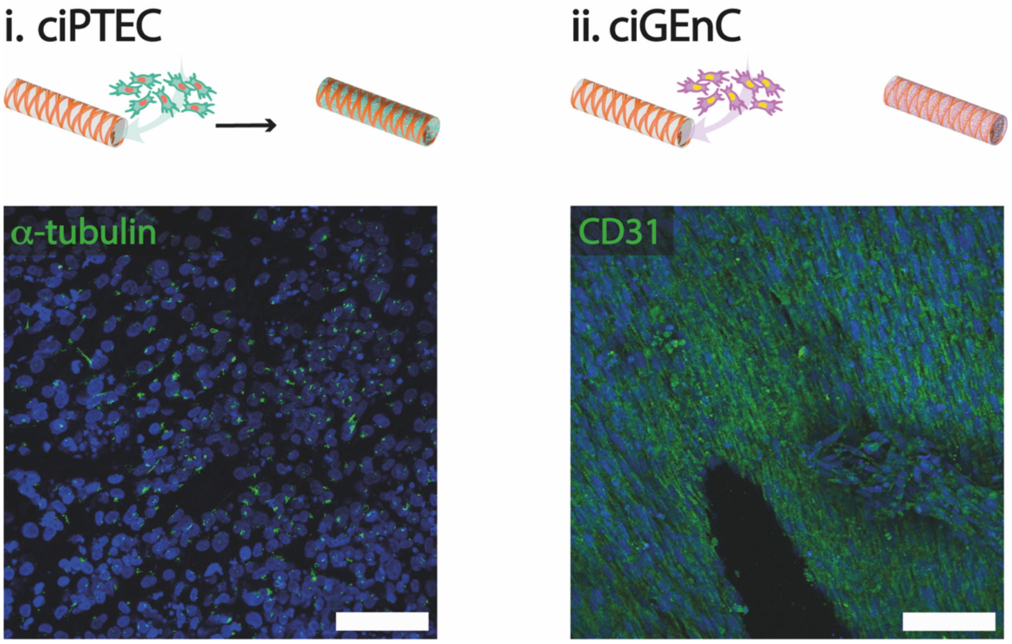


**Figure S10.** α-tubulin and CD31 stains in tubular (60° - 100 layers) scaffolds seeded with i. human conditionally immortalized proximal tubular epithelial cells (ciPTEC) and ii. glomerular endothelial cells (ciGEnC); scale bars: 100 μm.

| Primary antibody | Secondary antibody |
|---|---|
| Live / Dead™ Viability/Cytotoxicity Kit<br>L3224, Invitrogen | N.A. |
| AlexaFluor 488 phalloidin<br>1:1000<br>A22283 Thermo Fisher Scientific | N.A. |
| Rabbit polyclonal anti-collagen I | AlexaFluor 488 donkey-anti-rabbit |
| 1:100 | 1:200 |
| Ab34710, Abcam | Invitrogen |
| Goat monoclonal anti-collagen IV | AlexaFluor 568 donkey-anti-goat / AlexaFluor |
| 1:50 | 546 donkey-anti-goat / |
| 1340-01, Southern Biotech | 1:200 |
| 1340-01, Southern Biotech | 1:200<br>Invitrogen |
| Mouse monoclonal anti-α-tubulin | AlexaFluor 488 donkey-anti-mouse |
| 1:200 | 1:200 |
| T6793, Sigma-Aldrich | Invitrogen |
| Mouse anti CD31/PECAM1 | AlexaFluor 488 donkey-anti-mouse |
| 1:150 | 1:200 |
| BBA7, R&D Systems | Invitrogen |
| DAPI | N.A. |
| 1:1000 | |
| D9542, Sigma-Aldrich | |

**Table S1.** List of antibodies used for immunocytochemistry.
